# Characterisation of a *Teladorsagia circumcincta* glutathione transferase

**DOI:** 10.1101/2020.07.13.201368

**Authors:** Saleh Umair, Charlotte L.G. Bouchet, Qing Deng, Nikola Palevich, Heather V. Simpson

**Author notes:** Corresponding author Saleh Umair, AgResearch Ltd, Hopkirk Research Institute, Grasslands Research Centre, Tennent Drive, Private Bag 11008, Palmerston North 4442, New Zealand.

## Abstract

A 615 bp full length cDNA encoding a *Teladorsagia circumcincta* glutathione transferase (*Tc*GST) was cloned, expressed in *Escherichia coli* and the recombinant protein purified and its kinetic properties determined. The predicted protein consisted of 205 amino acids and was present as a single band of about 24 kDa on SDS-PAGE. Multiple alignments of the protein sequence of *Tc*GST with homologues from other helminths showed that the highest identity of 53-68% with haem-binding nematode proteins designated as members of the nu class of GSTs. Substrate binding sites and conserved regions were identified and were generally conserved. The predicted 3-dimensional structures of *Tc*GST and *Hc*GST revealed highly open binding cavities typical of this class of GST, considered to allow greater accessibility to diverse ligands compared with other classes of GST. At 25 °C, the optimum pH for *Tc*GST activity was pH 7, the V_max_ was 1535 ± 33 nmoles.min^-1^.mg^-1^ protein and the apparent K_m_ for the substrate 1-chloro-2,4-dinitrobenzene (CDNB) was 0.22 ± 0.01 mM (mean ± SD, n = 2). Antibodies in both serum and saliva from field-immune, but not nematode-naÏve, sheep, recognised recombinant *Tc*GST in enzyme-linked immunosorbent assays. The recognition of the recombinant protein by antibodies generated by exposure of sheep to the native enzyme indicates similar antigenicity of the two proteins. These findings could aid in the design of novel drugs and vaccine antigens for economically important parasites of livestock.

## 1. Introduction

Glutathione transferases (GSTs) (E.C. 2.5.1.18) are a large superfamily of enzymes, which have the principal function of protecting cells against oxidative stress, toxic, carcinogenic and mutagenic effects of endogenous substances and xenobiotics (Hayes and Pulford, 1995). The detoxification reactions involve the catalysis by GSTs of the conjugation of many electrophilic substances to the thiol group of the tripeptide glutathione (L-γ-glutamyl-L-cysteinylglycine) (Sheehan et al., 2001; Hayes et al., 2005; Deponte, 2013), followed by removal of the conjugated chemicals from the cells by transporters (Cole and Deeley, 2006). Additional functions of particular classes of GSTs include binding to hydrophobic molecules, modifying immune functions and participating in cellular metabolism and signalling (Brophy and Barrett, 1990; Board and Menon, 2013).

GSTs are universally present in bacteria and eukaryotes, in which multiple classes of the enzyme are expressed, although some classes have restricted distributions. The superfamily includes distantly related families of cytosolic GSTs (alpha, mu, omega, pi, sigma, theta and zeta), as well as mitochondrial/microsomal enzymes (kappa GSTs) and membrane-bound glutathione and eicosanoid metabolising enzymes (Hayes and Pulford, 1995; Sheehan et al., 2001; Hayes et al., 2005; Board and Menon, 2013). Proteins in the same cytosolic GST class have sequence identity of at least 40%, contrasting with less than 25% between classes (Oakley, 2011). The cytosolic enzymes are mainly responsible for detoxification, with the different classes showing a range of substrate affinities (Deponte, 2013). Other specific activities include immune modulation by the theta class of GSTs, which are MIF (macrophage migration inhibitory factor) protein homologues (Blocki et al., 1993), and the sigma GSTs, which have both pro- and anti-inflammatory functions in mammals and an immunomodulatory role in helminths (Flanagan and Smythe, 2011). The mitochondrial kappa GSTs are involved in energy and lipid metabolism (Petit et al., 2009; Morel and Aninat, 2011).

Many helminths express multiple GSTs, homologues of most classes of enzymes; these have been characterised using genomic and proteomic approaches (Brophy and Pritchard, 1994; Sheehan et al., 2001; Markov et al., 2015; Bae et al., 2016; Matoušová et al., 2016). Genome-wide sequencing is possible for some helminths and has allowed analysis of gene homology across the phylum (Campbell et al., 2001) and revealed the large number of genes encode for GSTs, e.g. around 50 different GST proteins in *Caenorhabditis elegans* (Markov et al., 2015). Detoxification of anthelmintic drugs by the numerous cytosolic GSTs is protective of internal parasites (Matoušová et al., 2016). MIF proteins (theta GSTs) have been identified in numerous species of helminth (Sparkes et al., 2017) and these proteins can modulate host immune responses to promote parasite survival (Matoušová et al., 2016). A family of haem-binding proteins, which also bind haematin, in the ruminant nematode *Haemonchus contortus* (van Rossum et al., 2004) and hookworms of the genera *Necator* and *Ancylostoma* (Zhan et al., 2005; Goud et al., 2012) have been assigned to the nu family, which may be a nematode-specific class, or possibly a subfamily of the sigma class (Markov et al., 2015).

Development of vaccines against parasitic helminths is an alternative control strategy to counter widespread anthelmintic resistance. Recombinant GST vaccines have provoked high levels of immune response and protection against cestode (Preyavichyapugdee et al., 2008) and hookworm infections (Zhan et al., 2005), suggesting GSTs could also be used in vaccines against other parasites. In the present study, the cDNA encoding a *Teladorsagia circumcincta* glutathione transferase (*Tc*GST) was cloned, expressed in *Escherichia coli* and the recombinant protein was produced and purified. *Tc*GST was verified as a GST protein by determining its kinetic properties in catalysing the conjugation of CDNB (1-chloro-2,4-dinitrobenzene) to the thiol group of L-glutathione. Enzyme-linked immunosorbent assays (ELISAs) were performed to determine if the recombinant protein was recognised by saliva and serum from sheep previously exposed to nematode parasites in the field.

## 2. Materials and methods

All chemicals were purchased from the Sigma Chemical Co. (Mo, USA) unless stated otherwise. Use of experimental animals for culturing and harvesting adult worms for RNA extraction has been approved by the AgResearch Grasslands Animal Ethics Committee (protocol #13052).

### 2.1 Parasites

Pure cultures of *T. circumcincta* were maintained in the laboratory by regular passage through sheep. Adult worms were recovered from the abomasa of infected sheep as described previously (Umair et al., 2013). Briefly, abomasal contents were mixed 2:1 with 3% agar and the solidified agar blocks incubated at 37°C in a saline bath. Clumps of parasites were collected from the saline soon after emergence and frozen in Eppendorff tubes at -80°C for molecular biology procedures.

### 2.2 RNA isolation and synthesis of cDNA

Adult *T. circumcincta* (50-100 µl packed volume) in 1 ml Trizol (Life Technologies) were ground to a fine powder in a mortar under liquid N_2_ and total RNA extracted according to the manufacturer’s instructions. The quality and concentration of the RNA was assessed, and first strand was synthesised from 1µg using the iScript Select cDNA Synthesis Kit (Bio-Rad) and a 1:1 mixture of Oligo (dT) _20_ and random primers. A full-length *T. circumcincta* GST sequence TDC00922-1 (AgResearch’s Internal database) was amplified from cDNA in a PCR containing the oligonucleotide primers *Tc*GST-FL-F (5’-ATCG<u>CATATG</u>GTTCACTACAGACTGCTT -3’) and *Tc*GST-FL-R (5’-CGAT<u>GCGGCCG</u>CGAATGGTGTGTTC -3’) and cloned into the expression vector AY2.4 (Knight et al., 2004), using the restriction enzymes Ndel and Notl (inserted into the forward and reverse primers, underlined in primer sequences, respectively) to allow the production of N-terminal His-tagged recombinant protein. The expression clone was sequenced to confirm the sequence identity.

Alignments were performed using the Muscle alignment option in Geneious Prime (Biomatters Ltd) with the Blosum 62 similarity matrix used to determine 100% similarity to a *H. contortus* amino acid sequence and other helminth GSTs. A second alignment against the Protein Data Bank (PDB) was carried out using the Position-Specific Iterative Basic Local Alignment Search Tool (PSI-BLAST) (Altschul et al. 1997).

### 2.3 Protein modelling and structural analysis of Tc*GST*

PSI-BLAST was used to compare the *Tc*GST and *Hc*GST protein sequences with deposited structures in the PDB. A structural model of *Tc*GST was constructed by submitting the amino acid sequence obtained to the I-TASSER server (Yang et al., 2015). For comparison, the amino acid sequence of the *H. contortus* GST (locus tag HCON_NP_LOC15789), located on chromosome 2 (GenBank accession number <u>CP035801</u>, BioProject accession number <u>PRJNA517503</u>) from the *H. contortus* NZ_Hco_NP genome v1.0 (Palevich et al., 2019a,b), was modelled and described as *H. contortus GST*. The structural model with highest C- and TM-score was further validated using Procheck (Laskowski et al., 1996) and ProSA-web (Wiederstein and Sippl, 2007). TM-score is a metric for measuring the similarity of two protein structures, or a global fold similarity between the generated model and the structure it was based on. Scores higher than 0.5 assumes the parent structure and modelled protein share the same fold while below 0.17 suggests a random nature to the produced model (Zhang and Skolnick, 2004). C-score is a confidence score for estimating the quality of predicted models by I-TASSER. It is calculated based on the significance of threading template alignments and the convergence parameters of the structure assembly simulations. C-score is typically in the range of -5 to 2, where a C-score of higher value signifies a model with a high confidence and vice-versa. The substrate binding domain was identified and active site residues were deduced and pictured using the PyMol molecular graphics system version 1.0 (Schrodinger).

### 2.4 Expression of T. circumcincta *recombinant* Tc*GST in* E. coli

*E. coli* strain BL21 (DE3) were transformed with *E. coli* AY2.4 *Tc*GST and grown in 10 ml Luria Broth (LB) supplemented with 100 µg/ml ampicillin for 16 h at 37 °C and 250 rpm. The culture was diluted 20-fold in LB with 100 µg/ml ampicillin and grown to OD_600_ 0.6-0.8 at 37 °C and 250 rpm. L-arabinose was added to a final concentration of 0.2% and the culture grown for an additional 3 h at 37 °C and 250 rpm. Bacteria were harvested by centrifugation at 5,000 *g* for 10 min at 4 °C. The pellet was weighed and the bacteria resuspended (10 g/ml) in equilibration buffer (20 mM sodium biphosphate, 0.5 M NaCl, 20 mM imidazole, pH 7.4). Protease inhibitors were added to the suspension, which was then passed through the chamber of a MP110 Microfluidizer® (Microfluidics, USA) seven times consecutively under ice at 20,000 psi to ensure the full lysis of *E*.*coli*, as recommended by the manufacturer. The crude lysate was centrifuged at 15,000 *g* for 20 min at 4 °C to remove cell debris and the supernatant filtered through a 0.22 µm filter prior to purification.

### 2.5 Purification of recombinant Tc*GST*

Purified recombinant polyhistidine protein was obtained by fast protein liquid chromatography (FPLC) under native conditions, using a Ni-NTA column (Qiagen), coupled to the Biologic DUO-FLOW BIO-RAD chromatography system (Bio-Rad, USA). Sodium biphosphate buffer was used as an equilibration buffer, sodium biphosphate containing 20 mM imidazole as the wash buffer, and sodium biphosphate containing 500 mM imidazole as elution buffer. The protein was dialysed overnight following the elution and the concentration was determined by the Nanodrop A280 nm assay, using the extinction coefficient 32890 M^-1^cm^-1^ and molecular weight 23.5 KDa.

### 2.6 Gel electrophoresis

SDS-PAGE was performed using NuPAGE Novex 4-12% Bis-Tris gels according to the instructions of the manufacturer (Life Technologies). Gels were stained with Coomassie Blue (Life Technologies). A western blot was performed on the purified protein, using a monoclonal anti-polyhistidine-peroxidase antibody. Blots were incubated overnight in 1:2000 antibody in buffer (4% skim milk powder in tris-buffered saline and 0.1% Tween-20) at room temperature and developed to detect His-tagged recombinant protein.

### 2.7 TcGST activity (E.*C. 2*.*5*.*1*.*18)*

*Tc*GST enzyme activity was measured at 25 °C by monitoring the conjugation of 1-chloro-2,4-dinitrobenzene (CDNB) to the thiol group of L-glutathione. The reaction product absorbs at 340 nm and the rate of increase in the absorption is directly proportional to *Tc*GST activity. The final reaction mixture (1 ml) contained assay buffer, enzyme mix, enzyme developer, recombinant protein (50 µg) and the substrate CDNB.

1. The optimum pH was determined over a pH range 6 to 9 with a substrate concentration of 5 mM glutathione and 1 mM CDNB. Subsequent assays were carried out at pH 7.
2. The apparent K_m_ for CDNB was determined in reaction mixtures containing 0-5 mM CDNB and 5 mM L-glutathione at pH 7.

### 2.8 ELISA

Pooled serum and saliva samples collected from parasite-naive and parasite-exposed sheep were tested by ELISA for the presence of antibodies that react with recombinant *Tc*GST. Serum and saliva samples were collected from 18 male 6-7 months-old Romney lambs previously exposed to multiple species of parasite, including *H. contortus* and *T. circumcincta*. These lambs had developed immunity against *T. circumcincta* infection. 5 µg/ml *Tc*GST were immobilised onto ELISA plates (Maxisorp, Thermofisher Scientific), free binding sites were blocked with Superblock (Thermofisher Scientific) followed by incubation for 2 h at room temperature with serial dilutions of serum (200- to 6400-fold) or saliva (20- to 160-fold) in ELISA buffer for 2 h at room temperature. Bound serum immunoglobulins were detected by incubation for 2 h at 37 °C with 1:4000 diluted rabbit anti-sheep IgG-HRP and colour development with 3,3’,5,5’-tetramethylbenzidine (TMB). Salivary IgA was similarly detected with rabbit anti-sheep IgA-HRP.

### 2.9. Data analysis

Replicate data are presented as mean ± SD. Graph Prism v5 was used to plot kinetic data and estimate K_m_ and V_max_.

## 3. Results

### 3.1 Tc GST gene sequence

The 615 bp full length *T. circumcincta* cDNA sequence, amplified from adult *T. circumcincta* cDNA, has been deposited in Genbank as Accession No. NX452942. Multiple alignments of the predicted *Tc*GST protein of 205 amino acids were made with helminth homologues, using Alignment Geneious 8 (Fig. 1). There was 53-68% identity with proteins from *Ancylostoma ceylanicum, Ancylostoma duodenale, H. contortus, Heligmosomoides polygyrus, Necator americanus, Oesophagostomum dentatum, Nippostronglus brasiliensis, Ancylostoma caninum, C. elegans* and *Caenorhabditis briggsae*. Identity was 26% or less with GST homologues from 9 other helminths. Substrate binding sites and conserved regions in other homologues were identified during protein modelling and are shown in Fig. 1.

**Fig. 1.**
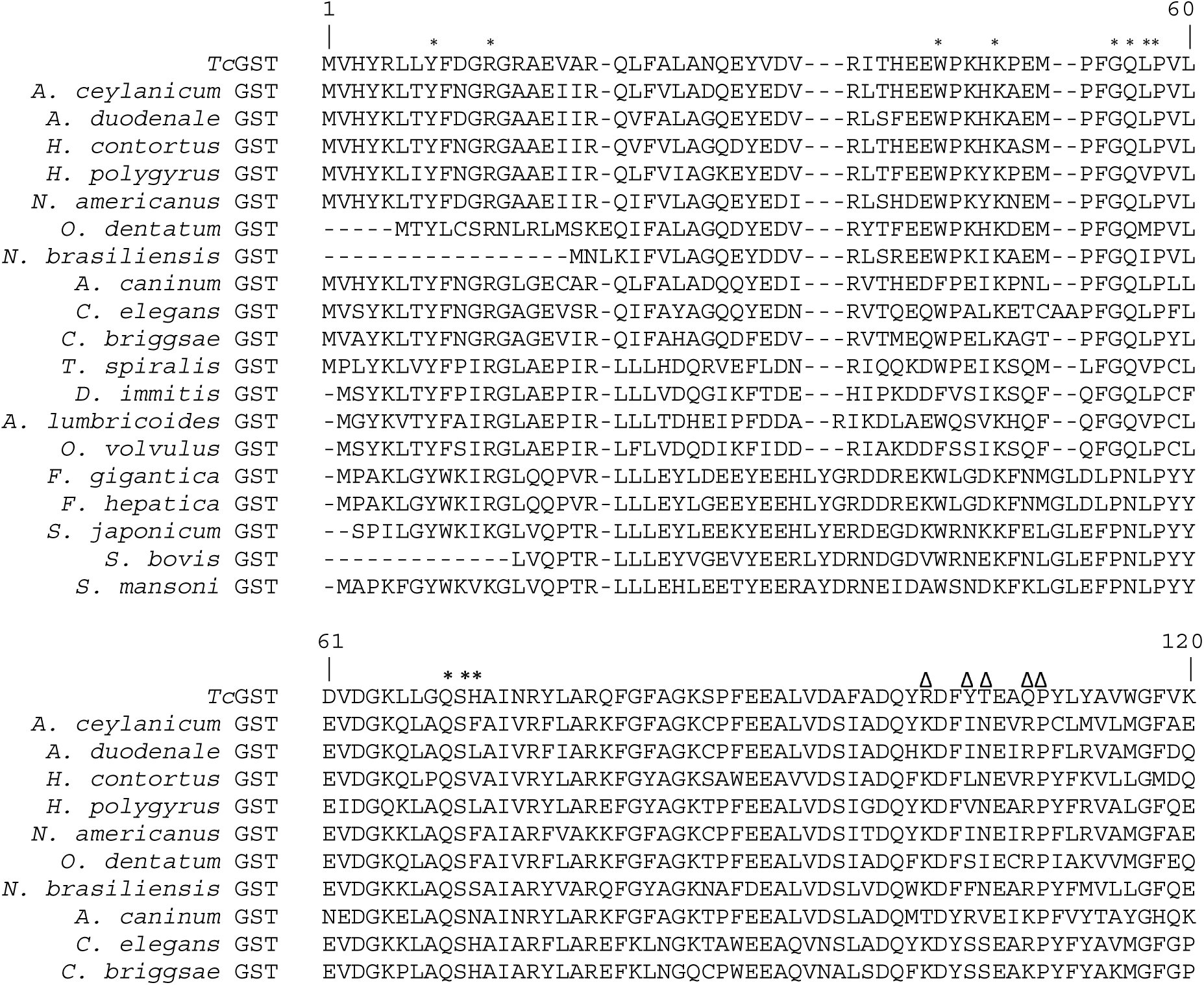
Multiple sequence alignment of *Tc*GST with homologues from *Ancylostoma ceylanicum* (EYC01088), *Ancylostoma duodenale* (GI: KIH60339), *Haemonchus contortus* (GI: AAF81283), *Heligmosomoides polygyrus* (GI: VDO85500), *Necator americanus* (GI: ACX53263), *Oesophagostomum dentatum* (GI: KHJ77903), *Nippostronglus brasiliensis* (GI: VDL81310), *Ancylostoma caninum* (GI: AAT37718), *Caenorhabditis elegans* (GI: CCD62662), *Caenorhabditis briggsae* (GI: XP002631478), *Trichinella spiralis* (GI: ABA42914), *Dirofilaria immitis* (GI: AAA21585), *Ascaris lumbricoides* (GI: ATZ35993), *Onchocerca volvulus* (GI: CAA54568), *Fasciola gigantica* (GI: ACH88355), *Fasciola hepatica* (GI: ADP09370), *Schistosoma japonicum* (GI: 62738608), *Schistosoma bovis* (GI: RTG90762) and *Schistosoma mansoni* (GI: AAA29888) homologues. The % identity of the helminth GST with that of *T. circumcincta* is shown at the end of the alignment. The triangles represent the non-specific substrate/chemical binding pocket, the H-site and those with an asterisk (*) represent the GSH-binding G-site. The % homology of each sequence with *Tc*GST is shown at the end of the alignment.

In the second alignment of the amino-acid sequence of *Tc*GST using PSI-BLAST, the protein sequence of *Tc*GST had the highest similarity (64.4%) to the *Hc*GST (HCON_NP_LOC15789) of *H. contortus* NZ_Hco_NP (Palevich et al., 2019a). This search also resulted in the assignment of a putative function to two of the top blast hits annotated as hypothetical protein (locus tag EYC01088 of *A. ceylanicum*) or proteins of unknown function (locus tag VDO85500 of *H. polygyrus*) with 68% and 63% identity respectively, based on the invertebrate non redundant (NR) database.

### 3.2 GST structure

The predicted 3D structures of *Tc*GST and *Hc*GST, and the binding and catalytic sites over a wide range of ligands are shown in Fig. 2. The protein structures for *Tc*GST and *Hc*GST were the superimposed best structural models corresponding to the monomer of <u>2ON5</u> (Asojo et al., 2007), associated with *Na-*GST-2 from the human hookworm *N. americanus*. The binding site and catalytic and active site residues that fall within 4 Å of the substrate (Tyr-8, Arg-14, Trp-39, Lys-43, Gly-49, Gln-50, Leu-51, Pro-52, Gln-63, Ser-64 and His-65) were similar in *Tc*GST and *Hc*GST (Fig. 2D). Both *Tc*GST and *Hc*GST had a TM Score of 0.90 ± 0.06, a root-mean-square deviation (RMSD) value of 2.8 ± 2.0 Å and normalized z-scores were less than 6.05. The main difference between the two structures was that *Tc*GST had a C-score of 1.33, whereas *Hc*GST had a C-score of 1.32.

**Fig. 2.**
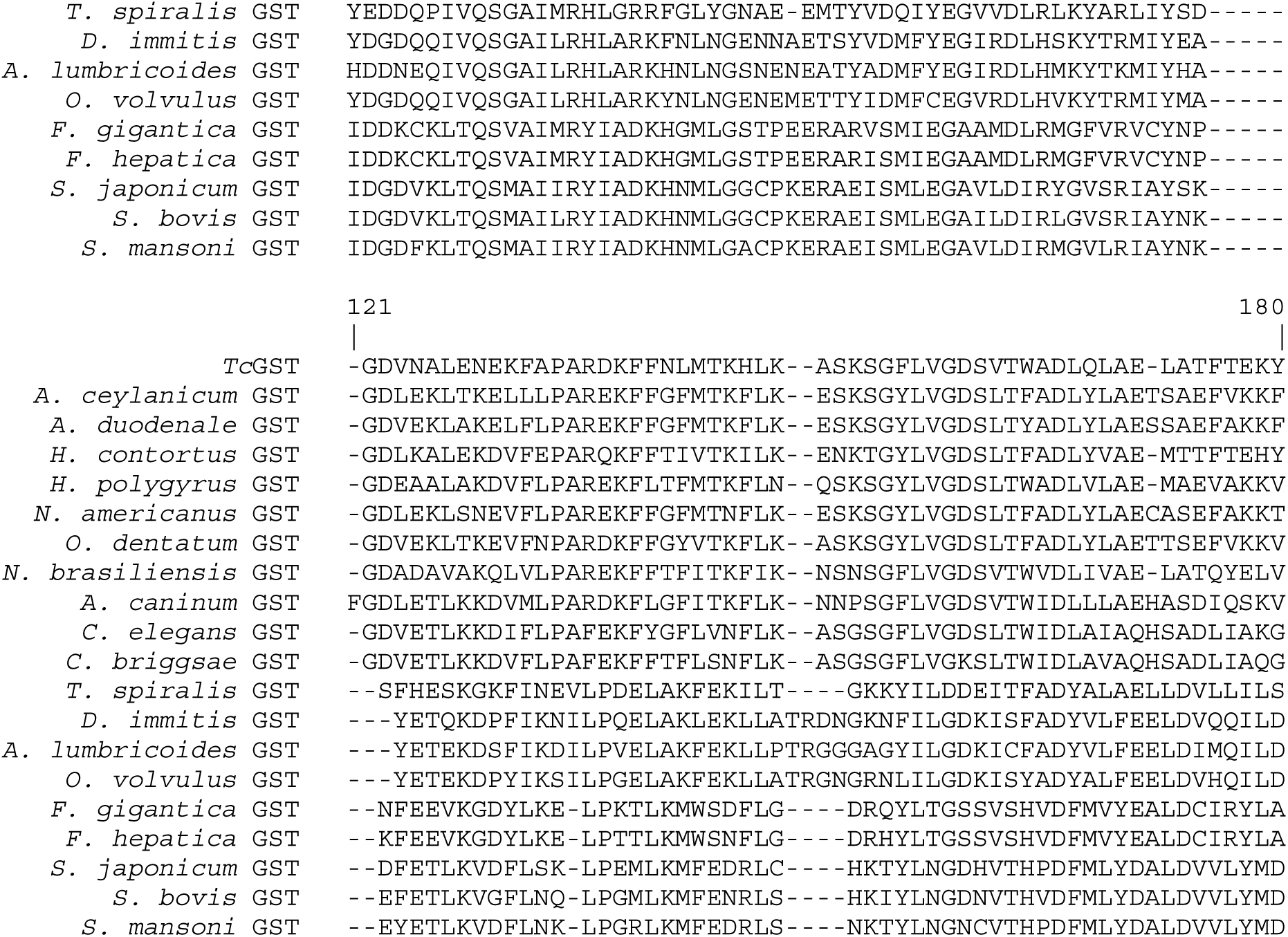
The predicted tertiary structure of the *Tc*GST and *Hc*GST monomers. (A) Location of the C- and N-termini in the predicted tertiary structure of *TcGST*. (B) Superposition of the predicted tertiary structure of *Tc*GST from *T. circumcincta* (red) and *H. contortus* GST (blue). (C) Location of the active site within *Tc*GST. (D) The active site of *Tc*GST (green) within 4Å of the superimposed 2CA8 (salmon) with polar bonds also shown in yellow.

### 3.3 Recombinant protein expression

Maximal production of functional recombinant GST was obtained in the *E. coli* strain BL21 (DE3) when expression was induced with 0.2% L-arabinose for 3 h at 37 °C. The purified N-terminal His recombinant *Tc*GST protein appeared as a single band of about 24 kDa on SDS-PAGE (Fig. 3A). The presence of a His-tagged recombinant protein was confirmed by Western blotting (Fig. 3B).

**Fig. 3.**
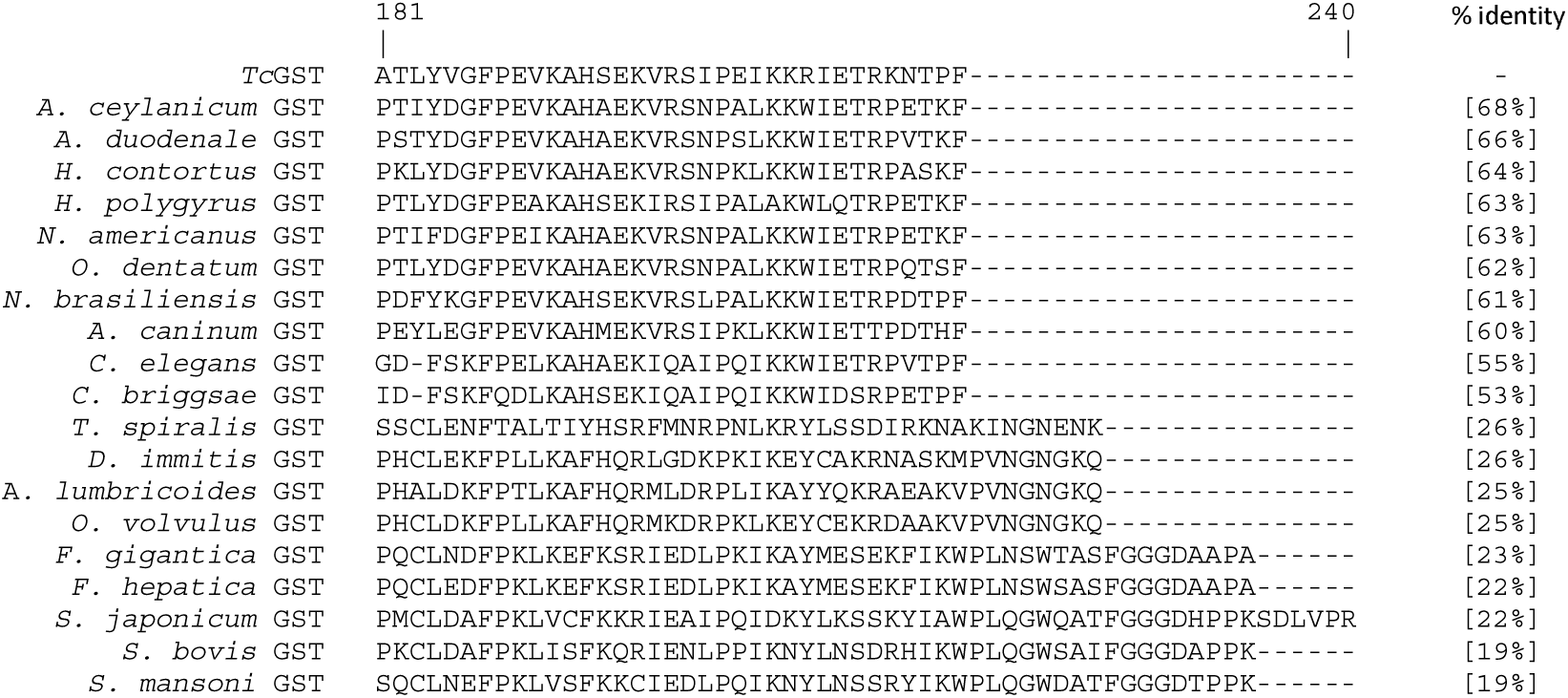
Purified recombinant *Tc*GST on a NuPage™ 4 - 12% Bis-Tris protein gel stained with SimplyBlue safe stain. Lane 1: Seeblue™ plus 2 Pre-stained protein standard in Kda; Lane 2: Filtered soluble bacterial lysate; Lane 3: Unbound material to HisTrap column; Lane 4: Elution fraction (purified recombinant *Tc*GST indicated by the black arrow).

### 3.4 Enzyme activity

The optimum pH for recombinant *Tc*GST activity at 25 °C was pH 7 (Fig. 4). The apparent K_m_ for CDNB was 0.22 ± 0.01 mM and the V_max_ 1535 ± 33 nmoles min^-1^ mg^-1^ protein (mean ± SD, n = 2) (Fig. 4). The Hill coefficient was calculated to be 1.70.

**Fig. 4.**
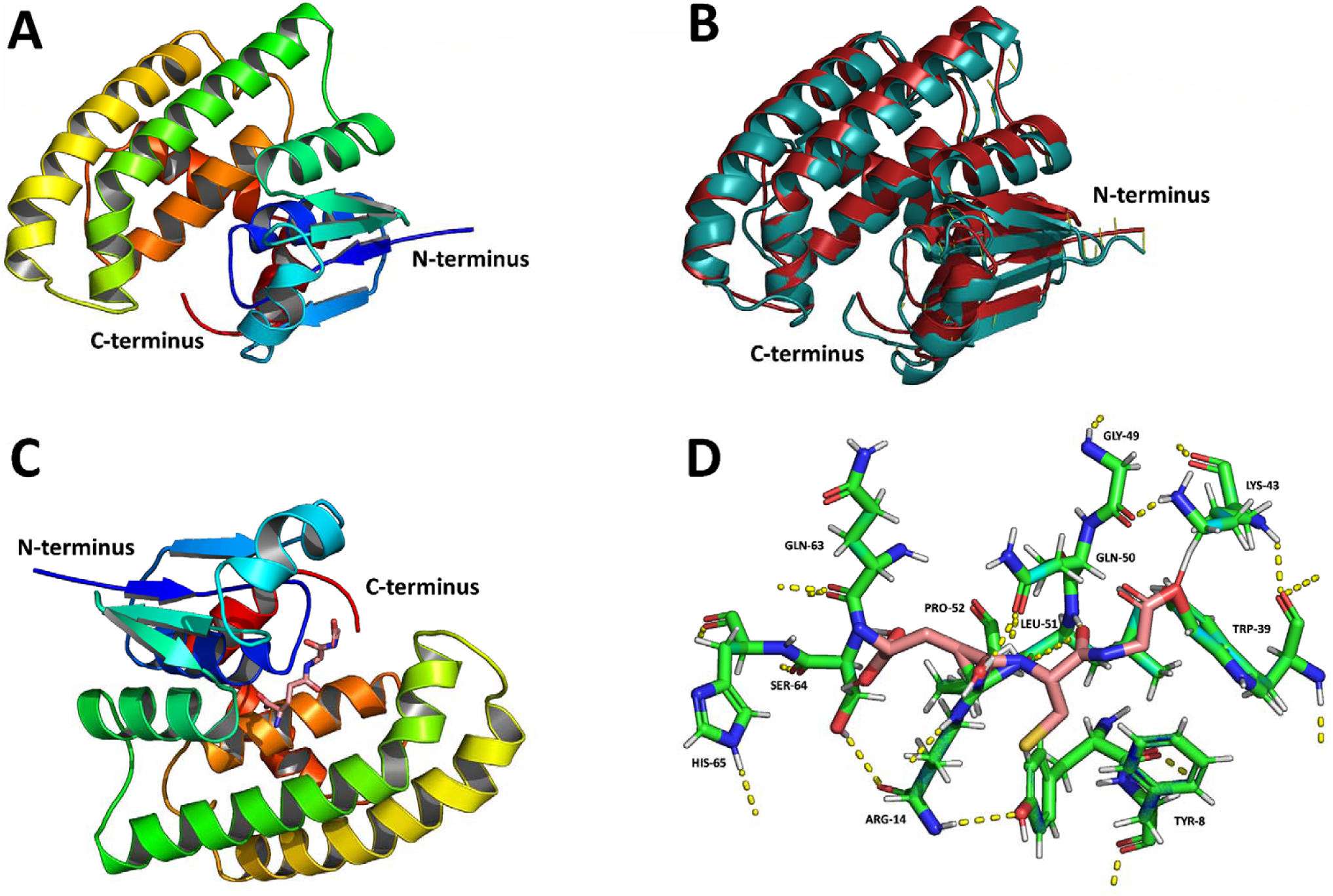
Effects of pH (top) and varying the substrate concentration at pH 7 (bottom) on the enzyme activity (mean ± SD, n = 2) of recombinant *Tc*GST at 25 °C. Activity was calculated from the conjugation of L-glutathione and 1-chloro-2,4-dinitrobenzene, monitored spectrophotometrically at 340 nm.

### 3.5 Host recognition

Recombinant *Tc*GST was recognised in an ELISA by antibodies in both serum and saliva collected from adult sheep exposed to nematodes in the field (Fig. 5). No antibody was detected when serum or saliva from parasite-naÏve animals was used.

**Fig. 5.**
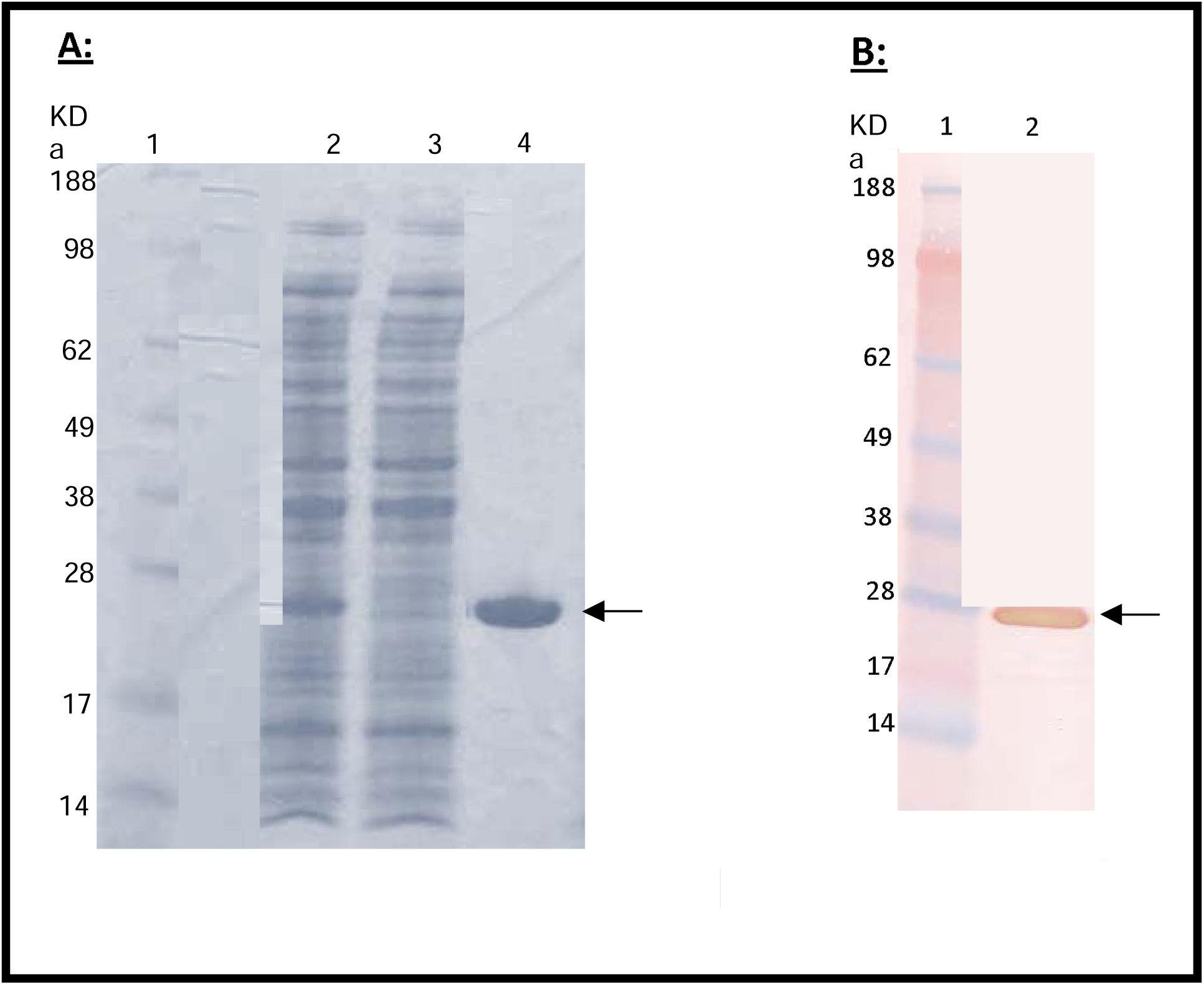
Recognition of recombinant *Tc*GST by serially diluted immune serum (IgG) (top) or saliva (IgA) (bottom) (■), but not by parasite-naÏve serum or saliva (•).

**Fig. 6.**
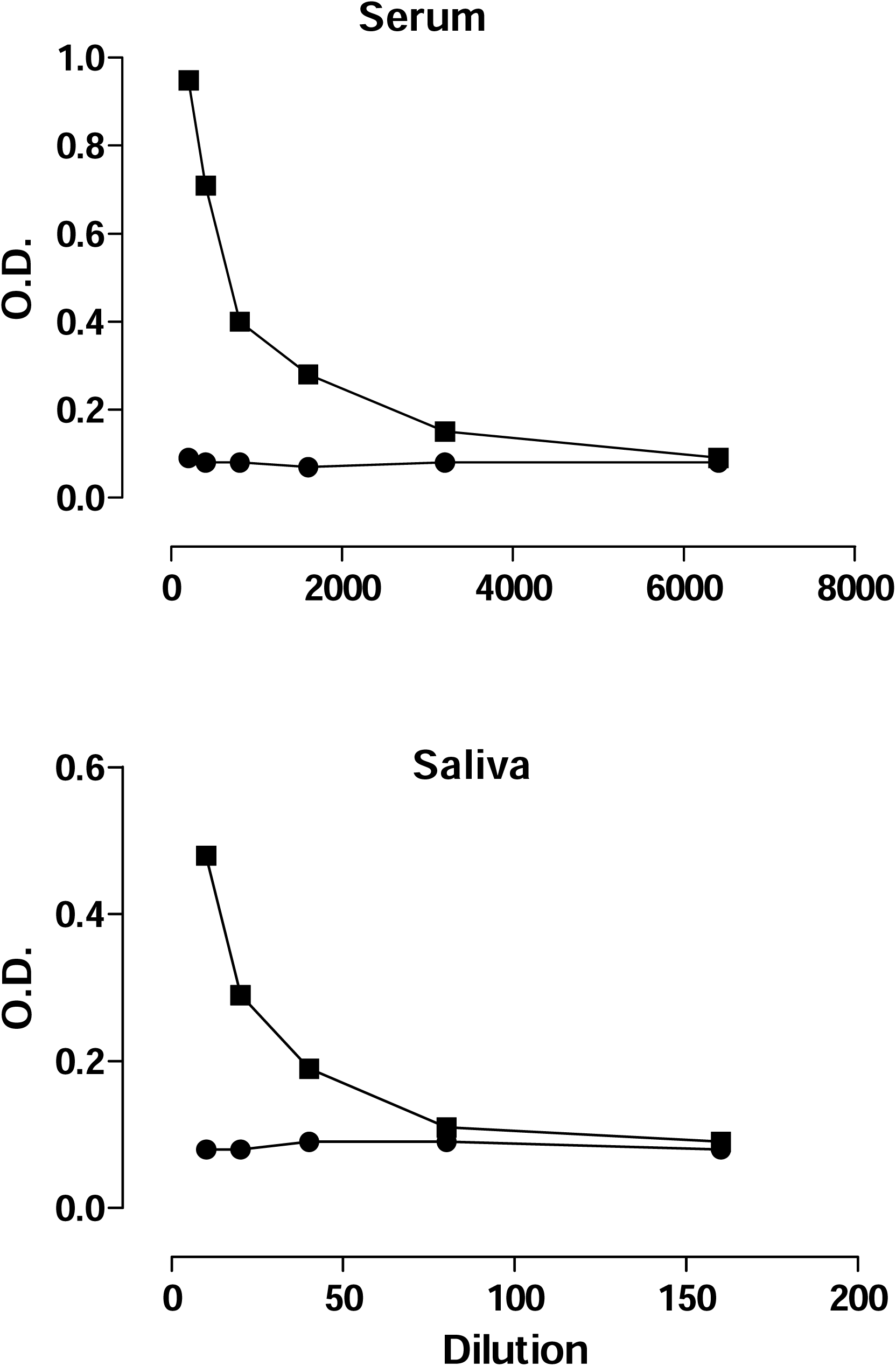
Recognition of recombinant TeciGST by serially diluted immune serum (IgG) (top) or saliva (IgA) (bottom) (⍰), but

## 4. Discussion

This study showed the close structural relationship between a *T. circumcincta* GST (*Tc*GST) and homologues from *H. contortus* and several animal parasitic nematodes and free-living species. This 615 bp full length cDNA sequence encoding *Tc*GST was amplified from adult *T. circumcincta* cDNA, cloned and expressed in *E. coli* and the 205 amino acid *Tc*GST protein was verified as a detoxifying enzyme capable of conjugating the substrate CDNB with L-glutathione. The protein was recognised by antibodies in both serum and saliva from field-immune sheep, but not nematode-naïve animals.

The GST superfamily is a large one, consisting of distantly related families of cytosolic and mitochondrial/microsomal enzymes, as well as membrane-bound glutathione and eicosanoid metabolising enzymes (Hayes and Pulford, 1995; Sheehan et al., 2001; Hayes et al., 2005; Board and Menon, 2013). The universal function of GST enzymes is detoxification of chemicals by catalysing their conjugation to the thiol group of glutathione before removal from the cell (Sheehan et al., 2001; Hayes et al., 2005; Cole and Deeley, 2006; Deponte, 2013). The *T. circumcincta* GST identified in this study is likely to be only one of the GSTs expressed in this species, as helminths are known to express homologues of most GST classes (Brophy and Pritchard, 1994; Sheehan et al., 2001; Markov et al., 2015; Bae et al., 2016; Matoušová et al., 2016); the genome of *C. elegans* contains around 50 different GST proteins (Markov et al., 2015). Database searches in the present study indicated that two of the top blast hits currently annotated as a hypothetical protein (locus tag EYC01088 of *A. ceylanicum*) and a proteins of unknown function (locus tag VDO85500 of *H. polygyrus*) can be assigned putative functions as GSTs. As more sequences become available for comparison, functions are likely to be progressively assigned to the large number (about 50%) of the protein-coding genes in helminth genomes of unknown function (Palevich et al., 2018).

*Tc*GST appears to belong to the nu GST class (Fig. 1), which may be a nematode-specific class, or possibly a subfamily of the sigma class (Markov et al., 2015), based on observations that proteins in the same GST class have sequence identity of at least 40%, contrasting with less than 25% between classes (Hayes et al., 2005; Oakley, 2011). These haem-binding proteins, which also bind haematin, have been characterised in the nematodes *H. contortus* (van Rossum et al., 2004), *Onchocerca volvulus* (Perbandt et al., 2005) and hookworms of the genera *Necator* and *Ancylostoma* (Zhan et al., 2005; Goud et al., 2012). Modelling the protein structures of *Tc*GST and *Hc*GST (Fig. 2) revealed that the best structural models corresponded to the monomer of <u>2ON5</u>, associated with *N. americanus Na-*GST-2 (Asojo et al., 2007). The searches of databases for other helminth GSTs allowed the assignment of a putative function as GSTs to two of the top blast hits currently annotated as a hypothetical protein (locus tag EYC01088 of *A. ceylanicum*) and a proteins of unknown function (locus tag VDO85500 of *H. polygyrus*). As more sequences become available for comparison, functions can become progressively assigned to the large number (about 50%) of the protein-coding genes in helminth genomes of unknown function (Palevich et al., 2018).

The universal function of GST enzymes is detoxification of chemicals by catalysing their conjugation to the thiol group of glutathione before removal from the cell (Sheehan et al., 2001; Hayes et al., 2005; Cole and Deeley, 2006; Deponte, 2013). Recombinant *Tc*GST conjugated the model substrate CDNB, with an optimum pH at 25 °C of pH 7 (Fig. 5), similar to those for nu class *H. contortus* and *A. caninum* GST, and with high activity (V_max_ 1535 nmoles.min^-1^.mg protein^-1^), similar to that of rHcGST-1 (van Rossum et al., 2004) and about twice that of Ac-GST-1 (Zhan et al., 2005).

GSTs have similar protein sequences (Fig. 1) containing a G-site, where glutathione binds, and the H-site, which is the non-specific substrate/chemical binding pocket where haem and haematin bind to nu class GSTs. These sites are shown in the proteins aligned in Fig. 1, where the triangles represent the largely conserved G-site and the asterisks the residues at the H-site. The molecular structures of nu class GSTs have been reported for HpolGSTN2-2 in *H. polygyrus* (Schuller et al., 2005), *Ov*-GSt2 in *O*.*volvulus* (Perbandt et al., 2005) and *Na*-GST-2 and *Na*-GST-2 in *N. americanus* (Asojo et al., 2007) and compared with closely related sigma class GSTs, such as *Na*-GST-3 in *N. americanus* (Kelleher et al., 2013). The best structural models of *Tc*GST and *Hc*GST (Fig. 2) corresponded to the monomer of *N. americanus Na-*GST-2 with similar catalytic and active site residues within 4 Å of the substrate at Tyr-8, Phe-9, Trp-39, Lys-43, Gln-50, Leu-51, Pro-52, Gln-63, Ser-64 and Val-65. Like other nu class GSTs, the overall binding cavities were more open and probably therefore more accessible to diverse ligands than other GSTs.

Nu class GSTs participate in nematode haem metabolism through their ability to bind both haem and haematin. Nematodes are unable to synthesise haem and require either an external source or a symbiont to supply haem, as well as transporters for its uptake across cell membranes and between tissues (Perally et al., 2008). Parasitic nematodes acquire haem from erythrocytes, host tissues and gut bacteria. A number of haem-responsive genes have been identified in nematodes, including the *C. elegans* transmembrane transporters *Ce-hrg-2*, expressed in the epidermis (Chen et al., 2012), and the *H. contortus* homolog *Hc-hrg-2*, which is expressed in all life cycle stages, but at the highest levels in L3 (Chen et al., 2012; Zhou et al., 2020). *A. ceylanicum Ace*-GST, a homolog of *Ac*-GST and *Na*-GST-1, is located in the epidermis, muscle and intestine of adult worms (Hang et al., 2020).

Recombinant helminth GSTs are showing promising results as vaccine antigens (Da Costa et al., 1999; Zhan et al., 2005, 2010; Preyavichyapugdee et al., 2008, Hang et al., 2020). Native *T. circumcincta* GST is highly antigenic and antibodies in both serum and saliva from field-immune sheep recognised recombinant *Tc*GST in an ELISA (Fig. 7), suggesting it also may be a useful antigen for inclusion in further studies to assess the protective efficacy of recombinant *Tc*GST in sheep and goats.

## Acknowledgments

The authors would like to thank Dr Jacqui Knight for the initial database search and Drs Sandeep Gupta and Sofia Khanum for critically reviewing the manuscript. The financial support of AGMARDT (Grant No. P14003) is gratefully acknowledged.

## Notes

### Competing Interest Statement

The authors have declared no competing interest.

